# A lipid atlas of human carotid atherosclerosis

**DOI:** 10.1101/2020.03.09.976043

**Authors:** AM Moerman, M Visscher, N Slijkhuis, K Van Gaalen, B Heijs, T Klein, P Burgers, YB De Rijke, HMM Van Beusekom, TM Luider, HJM Verhagen, AFW Van Der Steen, FJH Gijsen, K Van Der Heiden, G Van Soest

## Abstract

Carotid atherosclerosis is one of the main causes of stroke, mortality and disability worldwide. The disease is characterized by plaques, heterogeneous depositions of lipids and necrotic debris in the vascular wall, which grow gradually and may remain asymptomatic for decades. However, at some point a plaque can evolve to a high-risk plaque phenotype, which may trigger a cerebrovascular event. Lipids play a key role in the development and progression of atherosclerosis. Using matrix-assisted laser desorption/ionization mass spectrometry imaging, we visualized the distribution of approximately 200 lipids in 106 tissue sections of 12 human carotid atherosclerotic plaques. We performed unsupervised classification of the mass spectrometry dataset, as well as a histology-driven multivariate analysis. These data allowed us to compare the spatial lipid patterns with morphological plaque features that have been associated with enhanced risk of symptoms. The abundances of sphingomyelin and oxidized cholesteryl ester species were elevated specifically in necrotic intima areas, while diacylglycerols and triacylglycerols were spatially correlated to areas containing the coagulation protein fibrin. This is the first study to systematically investigate spatial lipid patterns in atherosclerosis, analyze their relation to histological tissue type, and to demonstrate a clear co-localization between plaque features and specific lipid classes and individual lipid molecules in high-risk atherosclerotic plaques.

## Introduction

Atherosclerosis, a lipid-driven inflammatory disease of arteries, is one of the main causes of death worldwide ^1^. The main driving mechanism for atherosclerotic plaque initiation is the gradual accumulation of lipids, originating from circulating lipoproteins, at sites of endothelial dysfunction in the vessel wall ^2^. The influx of lipids and their subsequent modification in the vessel wall triggers inflammatory reactions that exacerbate the atherogenic process ^3,4^. With disease progression, lipids start to play a dual role: they are deposited as metabolites of inflammatory processes but also form lipoproteins that act as signaling molecules, binding to a variety of macrophage-borne receptors ^5^. Previous studies have found that the lipid content of atherosclerotic plaques is different at various stages of disease progression ^6,7^. Thus, visualizing the molecular lipid composition of a plaque will give insight into local metabolic processes and reflect the stage of the disease ^8^.

Matrix-assisted laser desorption/ionization mass spectrometry imaging (MALDI-MSI) is a label-free molecular imaging technique and is suitable for detection and visualization of lipids in tissue sections ^9,10^. Systematic MSI studies of lipid distribution in human atherosclerotic plaques have not been performed to date, and only small numbers of samples have been studied experimentally using a limited spectrum of lipid species ^11–17^. In this study, we extend the range of atherosclerotic lipids of which spatial distributions have been mapped. We use a previously established MALDI-MSI pipeline for systematic imaging of lipids ^18^ to visualize the spatial distribution of 194 lipids in 106 tissue sections of 12 carotid atherosclerotic plaques, harvested at carotid endarterectomy surgeries in 12 patients. Analysis of this dataset identifies spatial lipid patterns in advanced atherosclerosis that transcend individual variability. We processed tissue sections adjacent to those studied by MSI for a series of histological staining procedures to identify compositional features of plaque vulnerability, i.e. necrotic core, the thrombus-associated protein fibrin, erythrocytes and foam cells^19–21^. In this way we were able to compare spatial lipid patterns with gold standard histological assessment of plaque composition. In this paper, we describe the systematic analysis of our highly dimensional MALDI-MSI dataset. Our initial approach was to assess the MALDI-MSI data for spatial correlations between lipids. Secondly, we considered the histological results and investigated correlations between lipids and compositional features of plaque vulnerability.

Thus, this unique study, in which the molecular lipid content of 12 human atherosclerotic plaques was systematically visualized with high resolution and contextualized in terms of gold standard histological plaque assessments, provides a first step towards identification of vulnerable plaque phenotypes based on a plaque’s lipid signature. Its findings might guide the development of imaging techniques that can detect the lipid profile of a plaque *in vivo* and drive the identification of plaques, and thus patients, in need of focal or systemic preventive treatment.

## Results

### MALDI-MSI of plaque lipids shows lipid class-specific spatial patterns

Twelve human carotid plaque specimens were harvested from patients undergoing carotid endarterectomy (CEA) surgery. Demographic information can be found in Table 1. A set of 10 μm thick axial cross-sections of the plaque specimens were collected at 2 mm intervals along the length of the plaque and collected on glass slides. One slide per cross-section was imaged using MALDI-MSI. This resulted in a total of 106 MALDI-MSI datasets. MALDI-MSI data was processed using a previously described workflow ^18^ and lipid images of 194 unique lipids were retained. A list of measured *m/z* values and the lipids that they represent is provided in Supplementary Table 2. Adjacent tissue sections were stained by a set of histochemical staining procedures, that enabled visualization of necrotic core (NC), the coagulation protein fibrin, foam cells (FC) and erythrocytes. Per tissue section, a segmentation image was made, in which these plaque components were delineated and their areas were measured. For all tissue sections per patient, Table 1 summarizes the average percentage area of histological components expressed per total intima area. This table demonstrates the heterogeneity in histological tissue composition between patients and within patients. More details on the histological compositions of the 106 individual tissue sections are provided in Supplementary Fig. 1.

**Table 1:** Average histological tissue compositions (percentages of intima area) per patient.

| Patient | Age [yrs] | Sex | Intima area<br>($\mu \pm SD$ (max)) [mm<sup>2</sup>] | % Necrotic core<br>($\mu \pm SD$) | % Foam cells<br>($\mu \pm SD$) | % Fibrin<br>($\mu \pm SD$) | % Erythrocytes ($\mu \pm SD$) | Number of sections | Representative tissue section* |
| --- | --- | --- | --- | --- | --- | --- | --- | --- | --- |
| A | 56 | F | 19.0 ± 10.4 (34.4) | 10.1 ± 9.0 | 1.1 ± 1.7 | 0.9 ± 1.9 | 0.0 ± 0.1 | 9 | |
| B | 69 | M | 22.2 ± 16.2 (54.5) | 14.4 ± 18.9 | 6.1 ± 7.8 | 10.3 ± 17.4 | 0.6 ± 2.1 | 12 | |
| C | 65 | M | 22.6 ± 6.9 (30.1) | 5.2 ± 4.8 | 5.1 ± 6.7 | 1.7 ± 3.5 | 0.0 ± 0.0 | 7 | 1 |
| D | 75 | M | 23.0 ± 13.9 (54.5) | 11.9 ± 17.6 | 4.6 ± 4.8 | 0.6 ± 1.1 | 3.7 ± 6.5 | 11 | |
| E | 75 | M | 25.4 ± 11.4 (47.2) | 8.5 ± 9.4 | 2.9 ± 1.8 | 14.4 ± 13.6 | 0.0 ± 0.1 | 8 | |
| F | 63 | M | 26.6 ± 11.6 (43.8) | 16.1 ± 11.0 | 3.1 ± 6.3 | 1.5 ± 2.3 | 0.3 ± 0.7 | 8 | 2 |
| G | 81 | M | 28.9 ± 17.2 (59.2) | 24.8 ± 17.1 | 0.6 ± 1.3 | 5.0 ± 5.9 | 0.0 ± 0.1 | 10 | |
| H | 62 | F | 35.7 ± 11.1 (51.0) | 8.7 ± 3.1 | 0.3 ± 0.4 | 5.6 ± 4.4 | 10.2 ± 7.7 | 5 | 3 |
| I | 69 | M | 49.6 ± 16.4 (70.1) | 48.6 ± 23.8 | 3.0 ± 5.1 | 15.7 ± 14.2 | 0.0 ± 0.1 | 7 | 4 |
| J | 79 | M | 52.1 ± 28.5 (101.0) | 35.3 ± 17.5 | 0.6 ± 1.1 | 23.6 ± 22.9 | 0.0 ± 0.0 | 8 | 5 |
| K | 69 | M | 53.3 ± 18.3 (77.4) | 47.5 ± 21.3 | 0.9 ± 1.4 | 13.8 ± 14.5 | 0.3 ± 1.1 | 12 | 6 |
| L | 82 | M | 57.3 ± 37.9 (120.0) | 25.6 ± 21.4 | 0.4 ± 0.7 | 32.0 ± 31.3 | 0.8 ± 1.4 | 9 | |

% area of histological component relative to total intima area. $\mu$ = mean, SD = standard deviation.
$\mu \pm SD$ : percentages were averaged over all tissue sections of a patient
$\mu \pm SD$ intima area is given in mm<sup>2</sup>, areas were averaged over all tissue sections of a patient. Intima area of largest tissue section is reported between brackets
\* Representative tissue sections are referred to in Figure 1, 2 and 4.

For a selection of lipids, Figure 1 illustrates the variety in spatial distributions over 6 tissue sections. The 6 tissue sections shown in this figure embody a representative subset of the 106 tissue sections, in terms of histological plaque composition. That is, all representative sections contained NC, four out of six sections contained fibrin areas of variable sizes (section 3, 4, 5, 6), section 2 contained a large FC area and in section 3 a large erythrocyte area was observed. The frequency of occurrence of different histologically-defined plaque components in the complete dataset is summarized in Supplementary Fig. 3M. The segmentation images of the representative tissue sections are shown in Figure 1.

**Figure 1:**
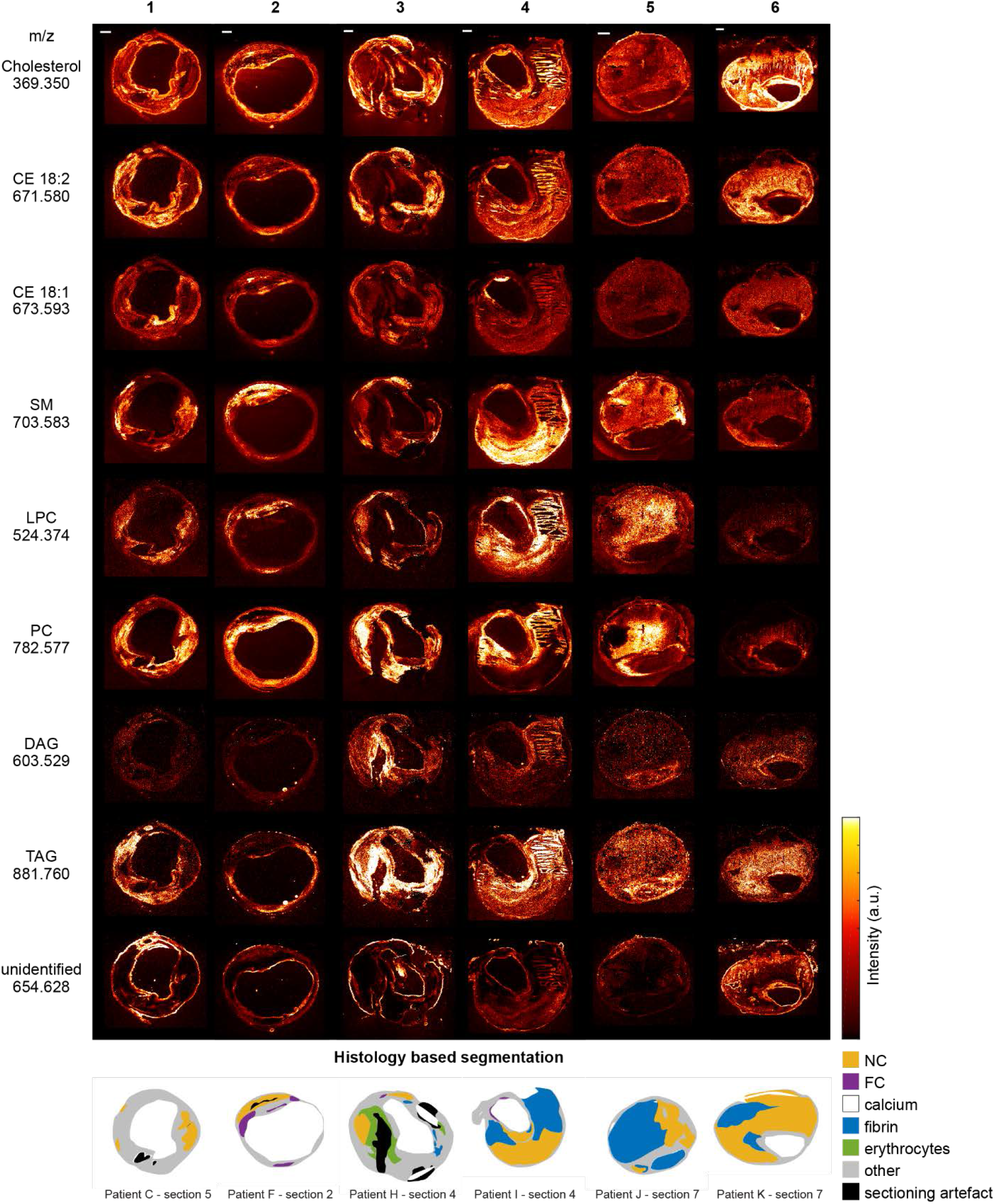
Overview of MALDI-MSI data for a selection of 9 lipids. Data is shown for 6 tissue sections representative of the 106 section dataset, in terms of histological plaque composition. For comparison, histology segmentation images are also shown. The lipid images were normalized per m/z value, to enable reliable comparison of the distribution of a specific lipid in different tissue sections. Also, CE(18:2) and CE(18:1) are shown on the same intensity scale, as are DAG and TAG. CE: cholesterol ester, SM: sphingomyelin, LPC: lysophosphocholine, PC: phosphocholine, DAG: diacylglycerol, TAG: triacylglycerol, NC: necrotic core, FC: foam cells. Scalebars are 1 mm.

In addition, we systematically compared the spatial distributions of a subset of 70 lipids, listed in Supplementary Table 2, for each tissue section by calculating the spatial cross-correlation between the lipid images. Per tissue section, the Pearson correlation coefficients between lipid images were visualized in a heatmap. The average heatmap of all tissue sections (Figure 2a) and the heatmaps of the 6 representative tissue sections (Figure 2b) are shown. In general, lipids belonging to the same lipid class showed similar, lipid class-specific spatial patterns, which were evidenced by positive Pearson correlation coefficients.

**Figure 2:**
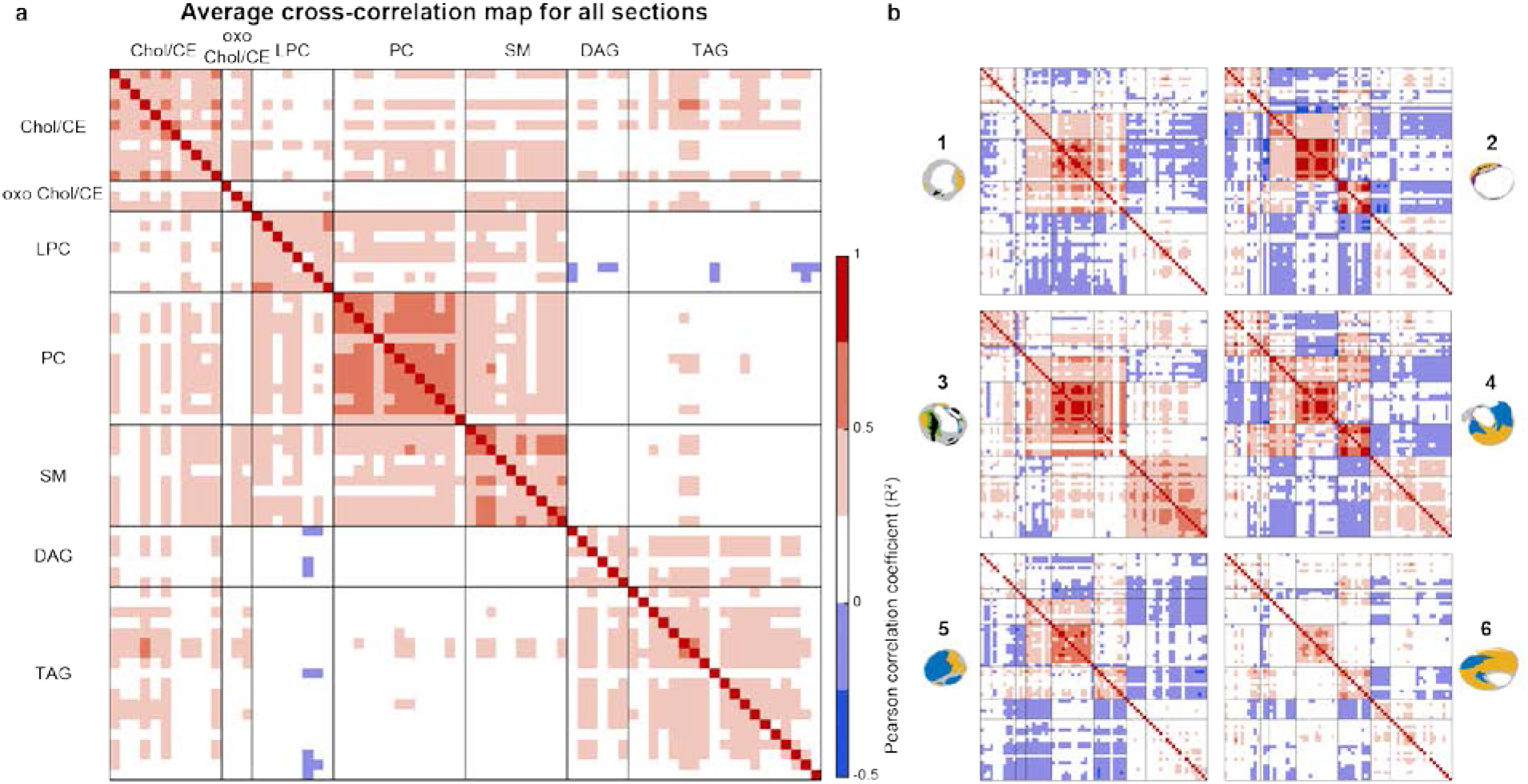
Spatial cross-correlation heatmaps. X- and Y-axes are the same and represent a selection of 70 m/z values (<u>Supplementary Table 2</u>), classified by lipid class. Per lipid class, the lipids are sorted from low to high m/z value. a) Average heatmap of all 106 tissue sections, showing a correlation within lipid classes and showing a moderate correlation between LPCs, PCs and SMs and also between DAGs and TAGs. b) Individual heatmaps of the 6 representative sections shown in Figure 1. Heatmaps of section 1 and 3 show moderate positive correlation between PC and SM lipids, whereas section 2 shows a negative between these lipids. CE: cholesterol ester, SM: sphingomyelin, LPC: lysophosphocholine, PC: phosphocholine, DAG: diacylglycerol, TAG: triacylglycerol.

As expected for atherosclerotic plaques, cholesterol was abundantly present in the mass spectra of all tissue sections in our dataset. Its presence was detected over the whole intimal area, though local variations in intensity were observed. No clear co-localization between cholesterol distribution patterns and histological components was seen (Figure 1). In general, cholesterol (*m/z* 369.351, [M-H_2_O+H]^+^) showed moderate cross-correlation with cholesterol esters (CEs) (Figure 2). The spatial patterns of two prominent CEs, CE linoleate (CE(18:2)) and CE oleate (CE(18:1)), show striking differences (Figure 1). While CE(18:2) is often present throughout the cross-section and in some cases strongly co-localizes with cholesterol, CE(18:1) is observed in high intensity spots around the lumen, while not abundant throughout the rest of the tissue sections (Figure 1). CE lipids in general showed mild spatial cross-correlation with cholesterol and with triacylglycerols (Figure 2).

A lipid with *m/z* 401.343, annotated as 7-ketocholesterol ^13^, was found in a subset of tissue sections. It showed a unique spatial distribution that was uncorrelated to the distribution of any other lipid in our dataset (Figure 2-oxChol first row). The spatial location of 7-ketocholesterol and cholesterol is depicted in Figure 3b for two tissue sections.

**Figure 3:**
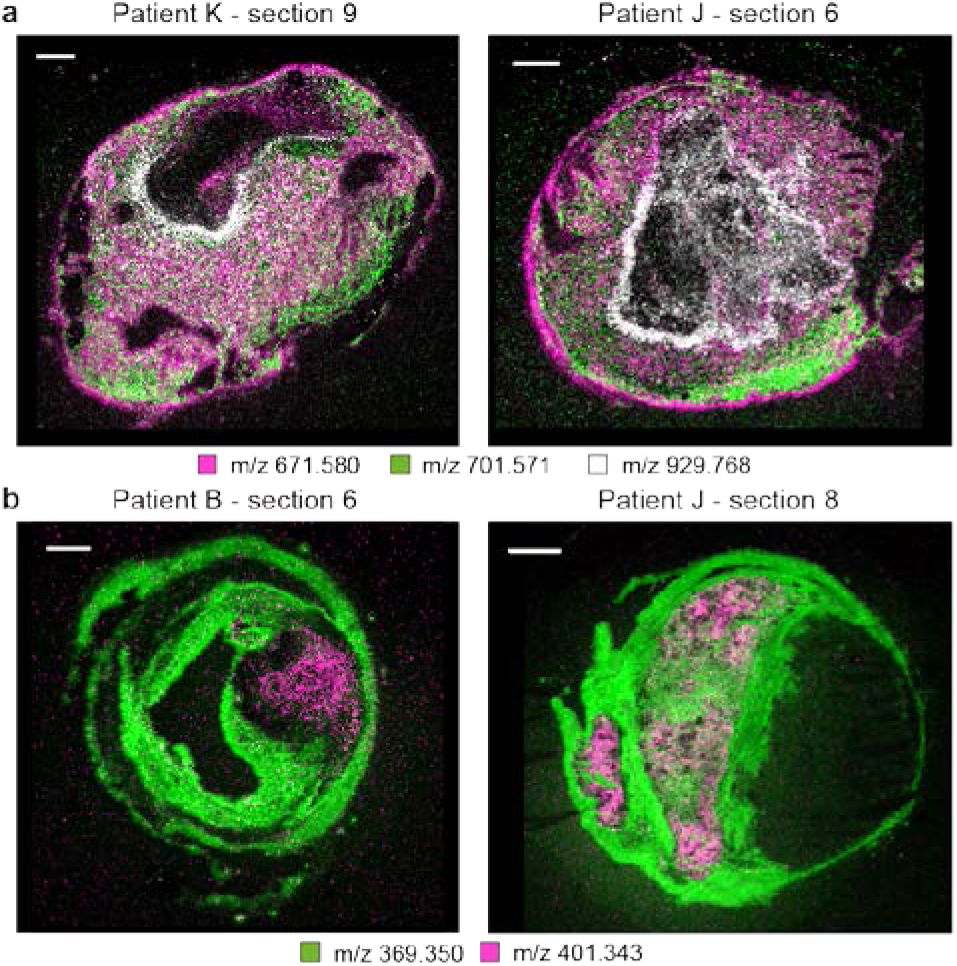
a) Combined MALDI-MSI image of spatial distributions of a CE in purple (m/z 671.580), SM in green (m/z 701.571) and a TAG in white (m/z 929.767) for three tissue sections. CE is found throughout the tissue section, while SM and TAG show a more localized spatial pattern. b) Spatial distribution of m/z 401.343, annotated as 7-ketocholesterol, in purple and m/z 369.350, identified as cholesterol, in green. 7-ketocholesterol shows a spatial distribution distinct from that of other lipids in our dataset. Scalebars are 1 mm.

We found that the co-localization of sphingomyelin (SM), lysophosphocholine (LPC) and phosphocholine (PC) lipids varied strongly: depending on the tissue section the spatial patterns of these three lipid classes could be similar but could also be completely different. In general, PCs were found to surround the lumen (Figure 1 – section 2, 6) and sometimes to exude into thrombotic or inflamed areas (Figure 1 – section 1, 3, 4, 5). The distribution of SMs did not deviate much of that of PCs in some tissue sections (Figure 1 – section 1) but in others SMs were clearly more abundant in the necrotic parts of the tissue section (Figure 1 – section 2, 4). These observations were quantified in the cross-correlation analysis: the degree of spatial correlation between SMs and PCs ranged from strong positive correlation, i.e. R^2^>0.75, to strong negative correlation, i.e. R^2^<−0.25 (Figure 2b – tissue section 2). In most tissue sections, the highest mutual spatial correlation was found between PC lipids (Figure 2). The distribution of PCs and LPCs was found to be weakly positively correlated in most tissue sections (Figure 2). LPCs are structurally similar to PCs but lack one fatty acid branch. When assessing the tissue sections that showed marked variations in PC and SM distributions (i.e. Figure 1 – section 2, 4, 5, 6) for the LPC distribution pattern, LPCs were highly expressed at the intersection of the locations that showed high SM and PC intensity. In contrast, in most tissue sections, the spatial patterns of the diacylglycerol (DAG) and triacylglycerol (TAG) lipids showed moderate to high correlation and were distinctive from the distributions of other lipids except for some CEs (Figure 2b - section 2, 4, 5, 6). However the intensities of measured DAG were much lower than those of TAG. The last column of Figure 1 represents MALDI images in which a high intensity signal is captured at the edges of the cross-sections, while the intensity in the rest of the section is negligible. We were not able to assign a lipid identity to this *m/z* value. Figure 3a shows overlay images of three lipids identified as a CE, a SM and a TAG for two tissue sections. The variation in spatial distributions between these lipids, which belong to different lipid classes, can clearly be seen.

### The spectral variety in the dataset can be captured by 6 spectral components

We reduced the dimensionality of the spectral data by unsupervised non-negative matrix factorization (NMF). In this way the spectral variety of the dataset was sparsely projected on a limited number of spectral components, which represented the major spectral patterns. We calculated that 6 was the optimal number of NMF components for our dataset. Figure 4a shows the spectra of the NMF components. Using this dimension reduction technique we were able to represent the 194 lipid images per tissue section in 6 images showing the relative intensities and distributions of the 6 spectral components. Component I consisted mostly of oxidized and non-oxidized cholesterol and CE lipids. Component II was a combination of cholesterol, both oxidized and non-oxidized, DAGs and a small proportion of LPCs. Component III was dominated by DAGs and TAGs. Component IV represented the unidentified *m/z* values. Component V consisted of LPCs, SMs and cholesterol, both oxidized and non-oxidized, while component VI represented the PC moiety of the data. Figure 4b shows the spatial distributions and relative intensities of the NMF components in the 6 representative tissue sections of our dataset.

**Figure 4:**
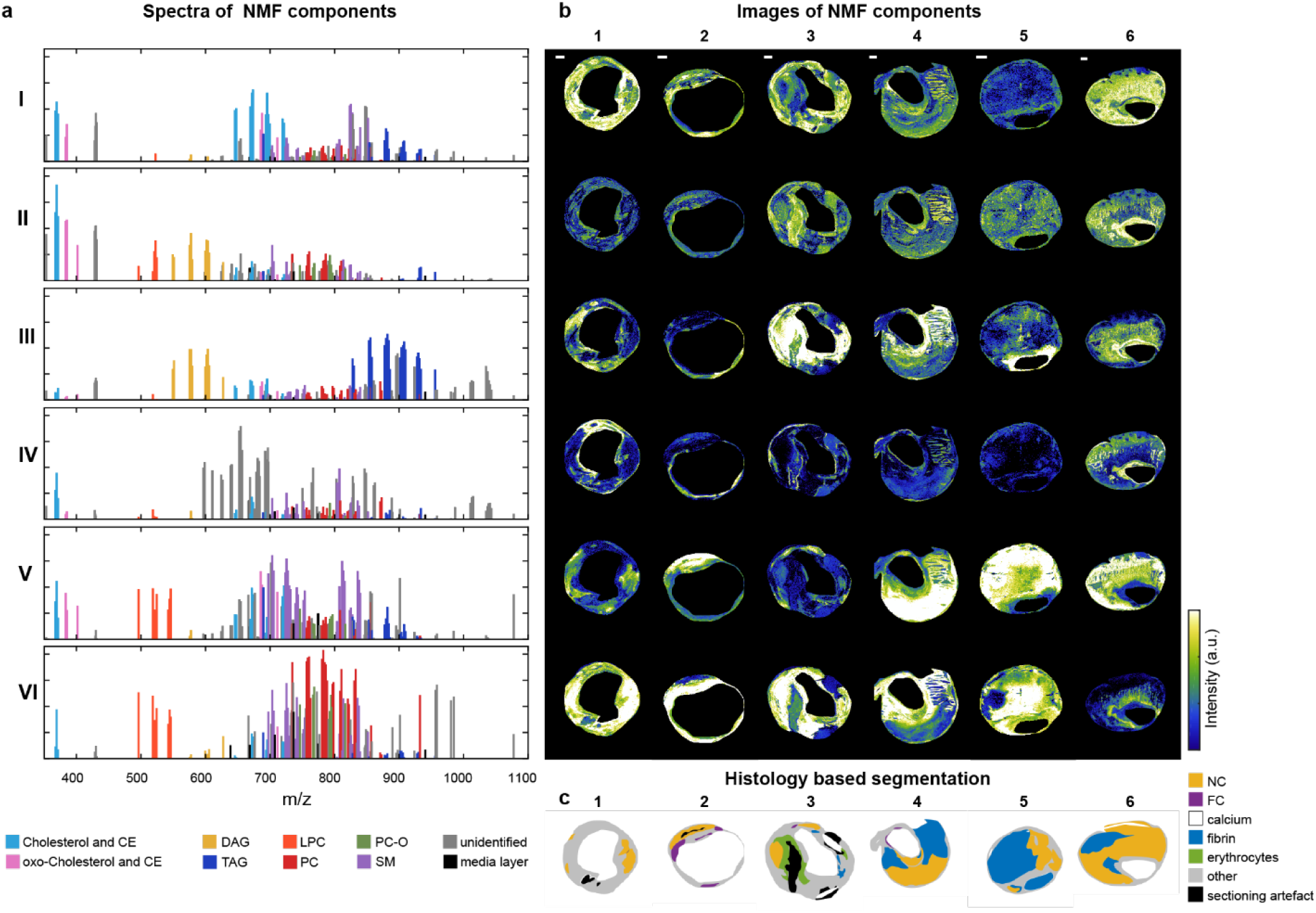
Unsupervised non-negative matrix factorization (NMF) of the 194 m/z-containing MALDI-MSI data. a) NMF spectra of the components showing the weight of each m/z value to that NMF component. m/z values in the NMF component spectra are labelled according to their assigned lipid class, showing a clear separation of lipid classes by means of NMF. b) Corresponding NMF-weighted images of the representative sections, showing the spatial distribution and relative intensities of the NMF components. c) Corresponding histology-based segmentation suggesting co-localisation between certain NMF components and histologically-relevant features. Scalebars are 1 mm. I-VI: NMF components, 1-6: representative tissue sections, CE: cholesterol ester, SM: sphingomyelin, LPC: lysophosphocholine, PC: phosphocholine, DAG: diacylglycerol, TAG: triacylglycerol, NC: necrotic core, FC: foam cells

### Correlation between spatial lipid patterns and histological plaque components

We registered the MALDI-MSI images and the histological segmentation images and compared the average spectra of different histological components by fitting a multivariate model. Four multivariate models, discriminating NC, fibrin, FCs and erythrocytes from all other tissue, were significant, i.e. R^2^ > 0.5 and Q^2^ > 0.5, for the combined data of at least 6 out of 12 patients. Model parameters of the significant models are listed in Supplementary Table 3. The multivariate models determine the *m/z* values that have the highest influence on separation between two histological components, reported as Variable Influence for Projection (VIP) values. VIP values > 1.0 are considered significant ^22^ and these are reported in Supplementary Table 4 for all significant models. These multivariate models found SMs and oxidized CEs to be more abundant in NC than in other intima tissue, while DAGs and TAGs were found to be distinctive for areas containing fibrin. The VIP values resulting from the FC model were mostly SMs, while the erythrocyte-related VIP values were mostly PCs.

In order to quantitatively establish the contrast in lipid composition between various tissues, we performed two univariate analyses of the intensities of *m/z* values with VIP > 1.0 appearing in the multivariate models: one contrasting NC and not-NC, and another contrasting fibrin and not-fibrin. Figure 5 shows the results of these univariate comparisons for NC versus not-NC tissue for two *m/z* values with significant VIPs. All *m/z* values that had significant VIP values in the multivariate model were significantly (p-value < 0.005) different between NC and not-NC in the univariate model, and the difference became stronger with increasing NC sizes (Figure 5b and 5c). The results of the univariate comparisons for two *m/z* values with VIP > 1.0 for fibrin versus not-fibrin are shown in Figure 6. In the univariate analysis of fibrin, 58 out of 62 *m/z* values in the VIP list were significantly different between fibrin and not-fibrin. The four non-significant *m/z* values had VIP values lower than 1.14.

**Figure 5:**
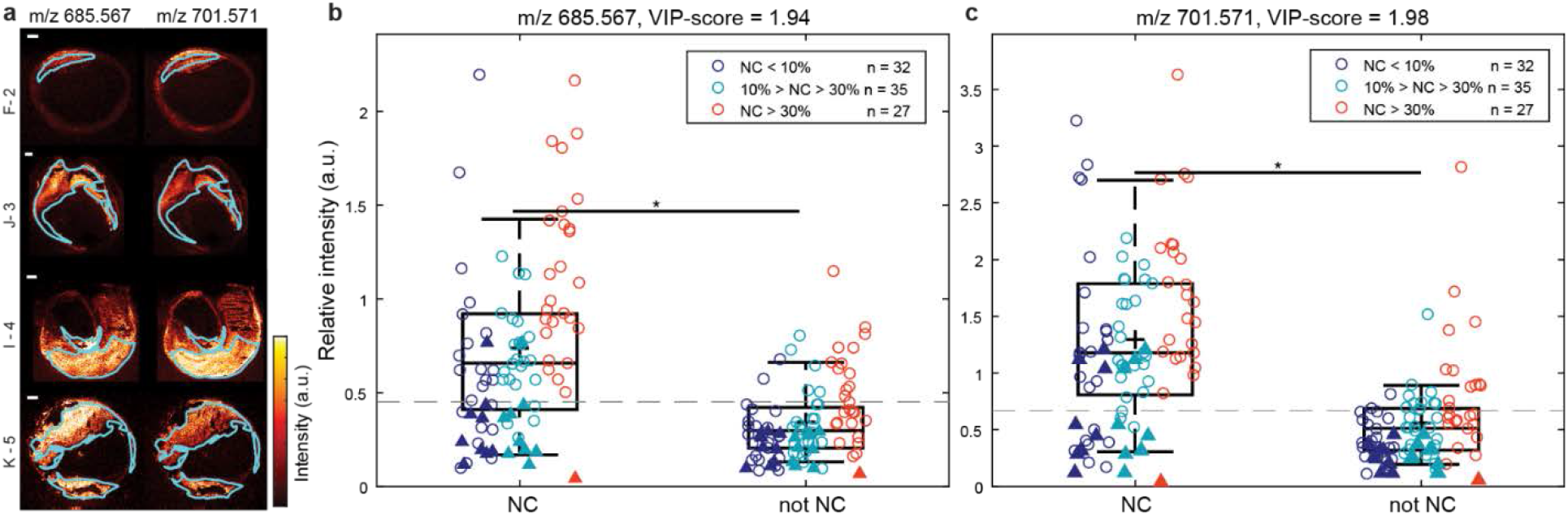
Univariate analysis of comparing intensities in necrotic core (NC) versus not-NC areas for two lipids (m/z 685.567 and m/z 701.571) with Variable Influence for Projection (VIP) values above 1.0. a) MALDI-MSI images of m/z 685.567 (oxoODE-CE [M+Na]^+^) and m/z 701.571 (SM(34:2) [M+H]+) of 4 different tissue sections with superimposed the outlines of the NC segmentation b) Boxplots of m/z 685.567 and c) m/z 701.571 showing the mean intensity of the pixels in the NC area and in the not-NC area. Datapoints are coloured according to the size of NC present in the corresponding tissue section. Only tissue sections containing NC were included in this analysis. The dotted line depicts the maximum Youden’s index, which represents the optimal cut-off intensity value in the tissue section above which the m/z value is more likely to be associated with NC. For m/z 685.567 the threshold is 0.45 and for m/z 701.571 it is 0.67. Both m/z values showed significantly different intensities between NC and not-NC (p-value = 1.8e-14 for m/z 685.567 and p-value = 2.5e-15 for m/z 701.571). o: data of the 9 patients that were included in the multivariate model, Δ: data of patients D, H and L, that did not fit the model. Scalebars are 1 mm.

**Figure 6:**
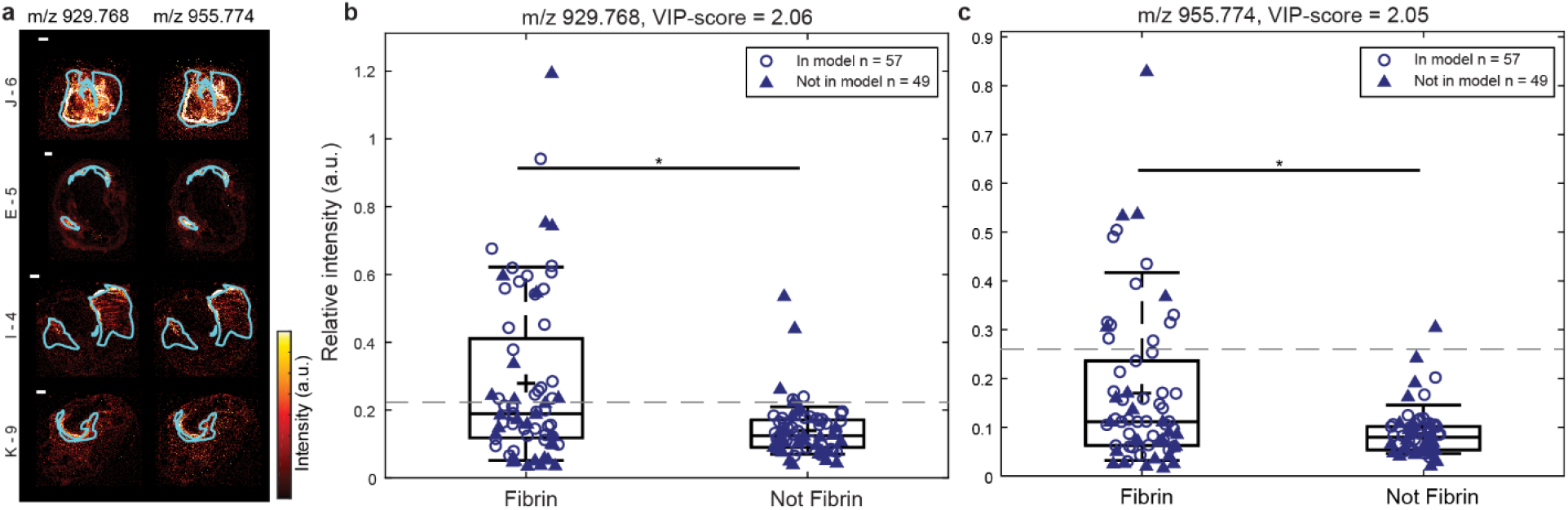
Univariate analysis of comparing intensities in fibrin versus not-fibrin areas for two lipids (m/z 929.768) and m/z 955.774) with Variable Influence for Projection (VIP) values above 1.0. a) MALDI-MSI images of m/z 929.768 (TAG(58:9) [M+H]^+^ or TAG(56:6) [M+Na]^+^) and m/z 955.774 (TAG(60:10) [M+H]^+^ or TAG(58:7) [M+Na]^+^) of 4 different tissue sections with superimposed the outlines of the fibrin segmentation b) Boxplots of m/z 929.768 and c) m/z 955.774 showing the mean intensity of the pixels in the fibrin area and in the not-fibrin area. Only tissue sections containing fibrin were included in this analysis. The dotted line depicts the maximum Youden’s index, which represents the optimal cut-off intensity value in the tissue section above which the m/z value is more likely to be associated with fibrin. For m/z 929.768 the threshold is 0.22 and for m/z 955.774 it is 0.26. Both m/z values showed significantly different intensities between fibrin and not-fibrin (p-value = 5.6e-07 for m/z 929.768 and p-value = 2.3e-06 for m/z 955.774). ○: data of the 6 patients B, C, F, G, J and K that were included in the multivariate model, Δ: data of patients A, D, E, H, I and L, that did not fit the model. Scalebars are 1 mm.

### Comparison NMF – multivariate analysis - histology

Since certain NMF components appeared to co-localize with areas identified in histology (Figure 4b and 4c), we investigated the similarities between the NMF spectra and the VIP values that were found to be distinctive for histological components. Per histological component, we investigated to which NMF component spectrum the *m/z* values with VIP > 1.0 were allocated. We defined an *m/z* value to be dominantly present in an NMF component spectrum when its intensity was higher than 0.4 times the normalized weight of that NMF component. As expected, the majority of the NC *m/z* values with VIP > 1.0 were found in NMF component V (Figure 7a). For fibrin, most VIP *m/z* values were found in NMF component III (Figure 7c). The majority of the significant VIP values for erythrocyte areas were most abundant in components VI and III and the VIP values associated with FCs corresponded to NMF component VI (Supplementary Table 5). Figure 7b and Figure 7d illustrate the correspondence between the histology-based multivariate analysis and the NMF analysis for NC and fibrin areas.

**Figure 7:**
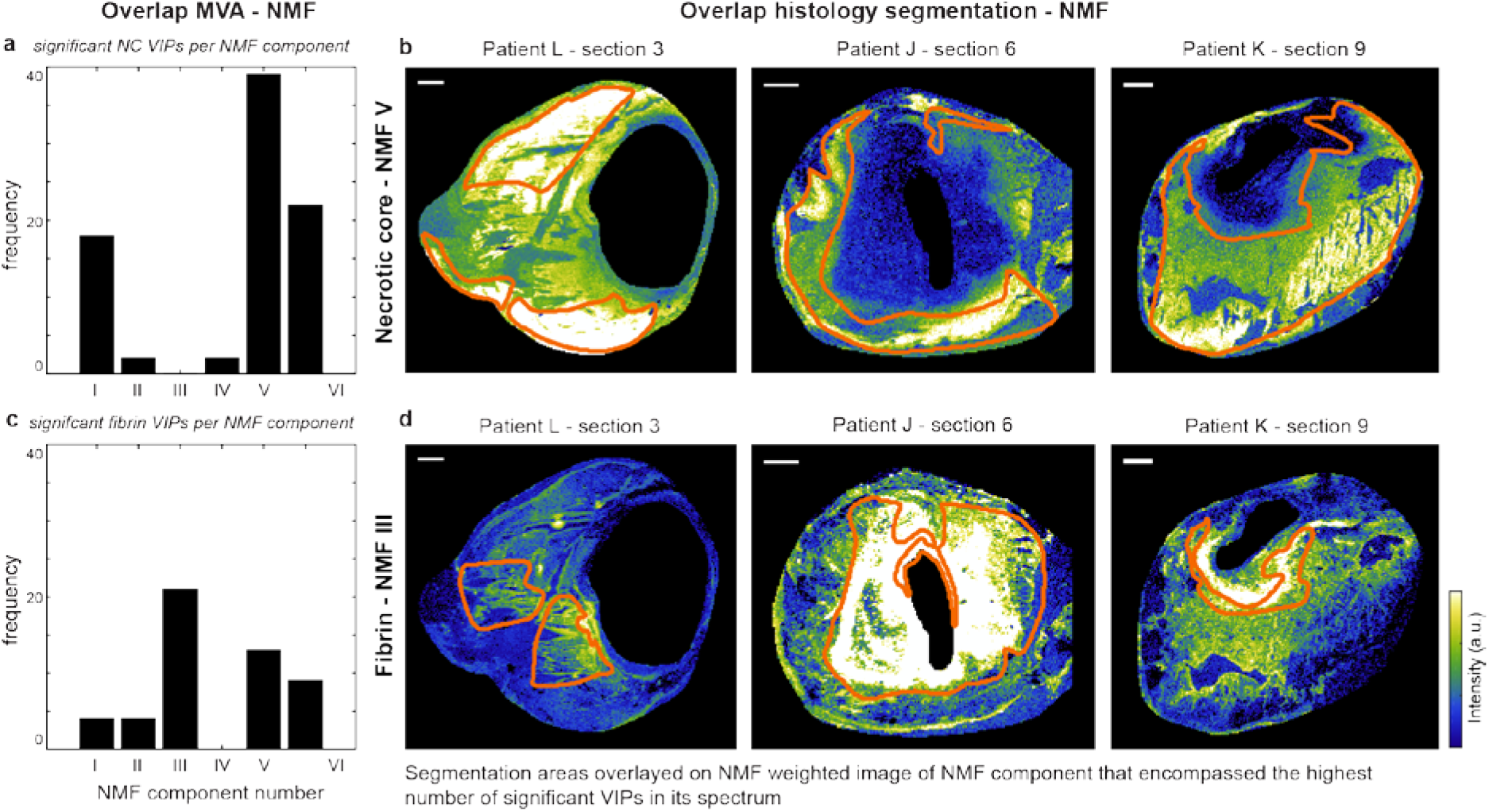
Overview of the correspondence between the histology-based multivariate analysis and the NMF analysis for NC- and fibrin-containing areas. a) Bar graphs showing the number of NC VIP values above 1.0 that were allocated to the different NMF component spectra. We considered an m/z value to be allocated to a specific NMF component when the intensity of this m/z value in the NMF spectrum was higher than 0.4 times the normalized weight of that NMF component. Most NC VIP values were allocated to NMF component V. b) The results in a are illustrated by depicting the NMF weight images for NMF component V for three tissue sections. c) Bar graphs showing the number of fibrin VIP values above 1.0 that were allocated to the different NMF component spectra. Most fibrin VIP values were present in NMF component III. d) The results in c are illustrated by depicting the NMF weight images for NMF component III for three tissue sections. Scalebars are 1 mm. MVA = multivariate analysis, NC: necrotic core, VIP: Variable Influence for Projection, NMF: non-negative matrix factorization.

## Discussion

With this study, we aimed to visualize the spatial distribution of lipids over the human carotid atherosclerotic plaque and to investigate the relation between lipids and compositional features of plaque vulnerability. We imaged the spatial distribution of 194 lipids in 106 tissue sections of carotid atherosclerotic plaques. This study collects the largest MSI dataset to date, spatially mapping the molecular lipid content of a series of human atherosclerotic plaques at high image and mass resolution, obtaining information on both the cross-sectional and the longitudinal composition of these plaques. Moreover, since we simultaneously processed adjacent tissue sections for histology, this study enabled the comparison of MALDI-MSI results with gold standard histological assessment of plaque stage.

In our dataset we observed lipids belonging to different lipid classes: cholesterol and CEs, LPCs, PCs, SMs, DAGs and TAGs. When stratifying the data by lipid class, we found that lipids belonging to the same lipid classes were distributed similarly over the plaque cross-sections, as quantified by their correlation coefficients. PC lipids in particular showed high mutual spatial correlation. PCs also showed spatial overlap with SMs and LPCs, while CEs were spatially correlated to DAGs and TAGs. However, the degree of spatial correlations was found to differ per tissue section. SMs and oxidized CEs co-localize in NC areas, and this co-localization became more apparent with greater NC size and thus with bigger intima area. The presence of oxidized CEs in NC is in line with theories on pathogenesis of atherosclerosis: during plaque progression, LDL proteins are thought be become oxidized and trapped inside the vessel wall ^2,23,24^. The presence of oxidized CE species in plaques, and their increase in abundance with lesion complexity, has been reported in literature ^11,25–27^. Increased SM levels in plaques, compared to healthy artery have also been described before ^14,28^, as well as the higher abundance of SMs, relative to PCs, with increasing plaque severity ^12,29^. Yet no previous study was able to visualize a clear co-localization of SM and oxidized CE lipid patterns with NC segmentations. We showed that the increased SM and oxidized CE levels concentrate in necrotic regions of the plaque, a finding that forms an important addition to the current understanding of the pathophysiology of atherosclerosis, and which also provides us a potential target for imaging NC presence or size in advanced plaques.

In literature, high levels of ceramides in serum, lipids that are structurally-related to SMs, have been associated with high-risk plaque ^30,31^. We detected low levels of ceramides in CEA tissue in the mass identification measurement using the Lipidyzer platform, but ceramide levels were probably under the detection limit of the MALDI-MSI experiments in this study or were suppressed by other ions.

In general, the spatial distribution of most cholesterol esters was similar to the distribution of cholesterol itself. Interestingly however, the spatial distribution of 7-ketocholesterol was distinct from that of other cholesterol-related molecules. In addition, this molecule showed a unique spatial distribution as it did not show any cross-correlation with other lipids in our dataset and was also not correlated with histological components. 7-Ketocholesterol has been found in atherosclerotic tissue before ^26,27,32^ and shown to play a pro-atherogenic role, e.g. by mediating cell death ^33,34^.

DAGs and TAGs showed a spatial distribution that was distinct from that of PCs, SMs and LPCs. Our multivariate analysis showed that a set of DAG and TAG lipids discriminated areas containing the thrombus-associated protein fibrin. A subset of TAG lipids was also associated with erythrocyte areas in the multivariate model. Previously ^18^ we described a co-localization between DAGs and thrombus in one of our tissue sections and this finding is thus confirmed in our current dataset. We hypothesize that the co-localization of DAGs and thrombotic plaque elements can be explained by the role that DAGs play in regulating protein kinase C (PKC), an enzyme known to induce an array of pro-atherogenic responses ^35^, including impairment of fibrinolysis ^36^ and platelet activation ^37,38^.

The tissue areas that contained FCs and erythrocytes were generally smaller than those of NC and fibrin. The lipids that were found to be distinctive for FC areas in the multivariate model were mostly PCs. When comparing the FC VIP values and the NMF component spectra, the majority of FC VIPs were present in the PC-dominated NMF component VI. Thus, there was large spatial overlap between the outcomes of the histology-based multivariate model and the NMF classification (Figure 7). A subset of erythrocyte-associated m/z values – as found by the multivariate analysis – were also PC lipids. PC lipids are major constituents of healthy cell membranes ^39^, which probably explains why the spatial distribution of PC lipids was found to correlate with areas of high cellularity, e.g. FC- and erythrocyte-rich areas.

Lipid identities could not be assigned to a subset of the 194 *m/z* values that were measured in the MALDI-MSI experiments. Most of the unknown masses in the MALDI-MSI data were not detected in the data captured by the Lipidyzer and FTICR measurements that were used for mass identification purposes. In addition, these unknown *m/z* values were not reported in the databases we consulted (Human Metabolome Database, SwissLipids and LipidMaps). The unidentified *m/z* values were dominating the spectrum of NMF component IV, while barely contributing to the spectra of the other NMF components. The distribution images of NMF component IV indicate that the unidentified *m/z* values were mostly present at the edges of the MALDI-MSI cross-sections. We assume that these images represent artefacts of the MALDI-MSI measurement method.

In our effort to identify lipids characteristic for plaque vulnerability, we conclude that certain SMs and oxidized CEs are significantly higher expressed in NC and that the abundance of these molecules is correlated to the size of the NC. Interesting observations to build further studies on are the correlation between DAGs and TAGs and thrombus fragments, i.e. erythrocytes and fibrin, providing a possible marker for intra-plaque bleeding. Additionally, erythrocytes and FCs are correlated with PCs. These findings add to the current understanding of atherosclerosis pathogenesis. In the future, these marker lipids may be used as targets for imaging and therapeutic applications.

## Methods

### Tissue collection and processing

Twelve human carotid endarterectomy (CEA) plaque specimens were surgically excised and were snap frozen and stored at −80°C until further processing. The surgery was performed using a protocol that preserves an intact lumen and plaque morphology ^40^. Upon processing, CEA specimens were divided in 2 mm thick cross-sections. Each cross-section was embedded in 10% porcine type A gelatin (Sigma-Aldrich, The Netherlands) and cryosectioned (CM3050 S, Leica Biosystems (cutting temps: OT − 21 °C; CT − 19 °C)) into 10 μm thick sections. Tissue sections were thaw-mounted on glass slides and stored at −80 ⁰C. One slide was processed for MALDI-MSI, 6 other slides were histochemically stained.

### Ethics statement

This study was performed according to the ethical guidelines sanctioned by the Ethics Board of Erasmus MC.

### MALDI-MSI sample preparation, measurement and data reduction

MALDI-MSI experiments and data processing were performed according to the methods described in Visscher et al ^18^. In short, we desiccated the tissue sections and applied 2,5-dihydroxybenzoic acid (DHB) matrix by sublimation (home-built sublimation system as described in Dekker et al. ^41^). MALDI-MSI experiments were performed on a Synapt G2Si-TOF system (Waters, Manchester, UK), operated in positive ion mode at the instrument’s resolution mode (single-pass reflectron TOF), using a 2000Hz Nd:YAG (355 nm) laser with a pixel-size of 45×45 μm^2^ and using Waters Research Enabled Software suite (WREnS). The mass range was 300-1,200 *m/z* and the laser fired 100 shots per pixel. The data was acquired using MassLynx v4.2 software (Waters, Manchester, UK), and HDI v1.4 was used to export the MSI data in imzML format and processed using an in-house developed data processing pipeline in MATLAB™ 2017a (Natick, USA) and mMass software ^42^. The processing pipeline consisted of the following procedures: (1) prepossessing, by smoothing and recalibration of the data using DHB cluster peaks, (2) peak picking form the base peak spectrum, (3) total-ion-current normalization and (4) removal of background *m/z* values and selection of lipid *m/z* values by a fractional mass filter and cross-correlations. We selected the lipid *m/z* values that were measured in more than 30% of all tissue sections and removed isotopes from the dataset, different adducts of the same molecule were retained. The data was log transformed for statistical analysis to comply with the assumptions of these methods^43^.

### Spatial cross-correlation

Per tissue section, the spatial correlation of lipid images was calculated using the Pearson correlation coefficient: 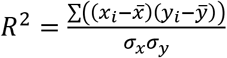. The results of all tissue sections were averaged to obtain the average Pearson correlation coefficients for our dataset. Results were visualized in heatmaps.

### Unsupervised clustering algorithm

We calculated the optimal number of spectral components for this dataset and we applied non-negative matrix factorization (NMF) dimension reduction to the combined spectral data of all tissue sections, using an NMF toolbox for biological datamining ^44–46^. We determined the optimum number of components based on dispersion coefficients ^47^.

### Histology, histology segmentation and image registration

The tissue section used in the MALDI-MSI experiment was stained for lipids by Oil Red O. Adjacent tissue sections were histochemically stained by: Miller’s elastic stain, Martius scarlet blue trichrome and hematoxylin-eosin. Based on the histological information, a segmentation image of the tissue section was drawn, in which we identified the following plaque components: necrotic core (NC), fibrin, foam cells (FC), erythrocytes and calcium. The segmentation images were registered to the MALDI tissue section by translation and scaling using an in-house developed point-based rigid image registration framework in MeVisLab (MeVis Medical Solutions AG, Germany), to enable correlation of histology and MALDI-MSI data. After registration, the mean spectrum for each plaque component was calculated per tissue section.

### Multivariate analysis

To investigate the presence of lipids characteristic for a plaque component, we performed multivariate analysis. For all plaque components, we fitted an Orthogonal Projections to Latent Structures Discriminant Analysis (OPLS-DA) model ^48^ in SIMCA 15 (Umetrics, Sweden) comparing the mean spectra of the plaque component to the mean spectra of the rest of the tissue. In these models the intensities of the *m/z* values in the mean spectrum were defined as variables and the histological components were defined as the observations. Quality of fit and predictability of the model were reported by R^2^ and Q^2^ values respectively. Significance of OPLS-DA models was checked by sevenfold cross-validation analysis of variance (CV-ANOVA) ^49^ and the models were validated by permutation tests. If the model was significant, we extracted the Variable Influence on Projection (VIP) values and defined *m/z* values with a VIP>1.0 to have a significantly different expression in the two histological components ^22^.

### Comparison unsupervised NMF clustering to histology-based multivariate analysis

Per NMF component spectrum we calculated a cut-off value of 0.4 times the normalized weight of that NMF component. We subsequently identified the *m/z* values with intensities above this cut-off value, resulting in a list of *m/z* values that dominated the NMF component. We compared these *m/z* values to the *m/z* values with VIP > 1.0 resulting from the multivariate analyses to check the correspondence between the NMF unsupervised clustering and the histology-based multivariate analysis.

### Statistics

The VIP values > 1.0 of the significant OPLS-DA models were separately tested on all data, including the patients for which the MVA model was not significant. Wilcoxon Signed Rank test was used to statistically test the non-log fold changes in intensity between the segmented and all other tissue areas. *M/z* values that had a p-value ≤ 0.05 were considered statistically significant. Receiver operating characteristic (ROC) was used to determine the optimal cut-off intensity (maximum Youden’s index) for the VIP *m/z* values.

### Lipid identification

We performed lipid identification experiments in two ways. Firstly we performed a lipidomics analysis on homogenized carotid endarterectomy tissue using the Lipidyzer platform (Sciex, Framingham, MA). In short, we divided atherosclerotic plaque tissue (~ 50 mg) originating from various locations of the CEA sample over nine vessels. The tissue was homogenized in 2 mL ice-cold methanol/MilliQ water (50/50 % (v/v)) with three 10 s bursts using an ultraturrax homogenizer. Homogenates were transferred to 2 mL Eppendorf tubes and stored on ice until centrifugation at 20,000 × *g*. The supernatant was collected and submitted to lipidomics analysis. Samples were processed according to manufacturer protocol before MS/MS analysis. Briefly, 50 μL of tissue homogenate was mixed with the software pre-calculated amount of isotopically-labelled internal standard compounds in 13 lipid classes before liquid-liquid extraction with dichloromethane (DCM)/methanol. After 30 minutes, the lower phase was collected and the sample re-extracted with DCM. The combined extracts were dried under flowing N_2_ and re-suspended in running buffer (10 mM ammonium acetate in DCM/methanol 50/50 (% v/v)).

The extracted samples were analyzed using the manufacturer-provided Lipidyzer SRM methods on a Shimadzu Nexera 20 UPLC system hyphenated to a Sciex 5500 Q-TRAP mass spectrometer equipped with a Selexion differential ion mobility module run with 1-propanol as a carrier. After analysis, lipid species quantification was performed using the Lipidyzer software.

Additionally, we performed MALDI-FTICR measurements on two plaque tissue sections with large intima area. MALDI-FTICR-MSI was performed on a 12 T Bruker Daltonics solariX xR mass spectrometer equipped with dynamically harmonized ParaCell™, and a CombiSource™ (Bruker Daltonics, Bremen, Germany). The instrument was operated using ftmsControl (v2.1.0 Build 98; Bruker Daltonics), and data were collected using a transient length of 0.4194 s (512k data point time domain), resulting in an estimated resolution of 97,000 at *m/z* 400. MALDI-measurements were performed using the SmartBeam™-II laser (λ = 355 nm) operating at 200 Hz, at 21% power, and using the “Small” focusing setting (ablation area approximately 70×70 μm^2^). A total of 50 shots were acquired per pixel, with a mass range covering 294.86-1,500 *m/z*. MSI analyses were performed at 70×70 μm^2^ and 100×100 μm^2^ pixel-size. Acquired data was loaded in SCiLS Lab (v2016b; Bruker Daltonics), and converted to imzML before further processing. We uploaded the FTICR results to the METASPACE annotation platform ^50^ and we searched the available annotation databases with a false-discovery rate (FDR) < 10%.

## Acknowledgements

We would like to thank Aad van der Lugt for helping us with clinical sample collection. This project was funded by ZonMw—project number: 91113020, Dutch Heart Foundation—project number: NHS2014T096 and Nederlandse Organisatie voor Wetenschappelijk Onderzoek—project number: 16131

## Author Contributions

A.M.M. and M.V. collected tissue, performed MALDI-MSI experiments, analyzed data and wrote the manuscript; N.S. performed multivariate analysis; K.v.G. performed histological analyses; B.H., T.K. and Y.B.d.R. performed lipid identification experiments; P.B., T.M.L. and H.M.M.v.B. supported methods development; H.J.M.V. assisted with clinical sample collection; K.v.d.H. and G.v.S. initiated the research, designed the study, and together with A.F.W.v.d.S. and F.J.H.G. supervised the project. All authors critically reviewed the manuscript.

## Competing Interests statement

The authors declare no competing interests.

## Materials & Correspondence

Supplementary information for this paper is available online. Correspondence and requests for materials should be addressed to G.v.S.

