## Supplementary information for "A lipid atlas of human carotid atherosclerosis"


| **Content** | | **Page** |
|---|---|---|
| Supplementary Table 1 | List of 194 lipids measured in human carotid atherosclerotic plaques | 2 |
| Supplementary Figure 1 | Histological tissue composition of 12 carotid plaques | 5 |
| Supplementary Table 2 | OPLS-DA model parameters | 6 |
| Supplementary Table 3 | Lists of *m/z* values with VIP>1.0 resulting from significant OPLS-DA models | 7 |
| Supplementary Table 4 | Number of *m/z* values with VIP > 1.0 for the different multivariate models | 11 |
| Supplementary References | | 11 |

Supplementary Table 1: List of 194 lipids measured in human carotid atherosclerotic plaques

| ***m/z ± 0.02*** | **lipid class** | **ID** | **adduct** | ***m/z ± 0.02*** | **lipid class** | **ID** | **adduct** |
|---|---|---|---|---|---|---|---|
| 353.331 | unknown | | | 638.472 | unknown | | |
| 367.343 | Chol | Cholesterol derivative[LM,d] | [M-H2O+H]+ | 638.587 | unknown | | |
| 369.350* | Chol | Cholesterol[c] | [M-H2O+H]+ | 640.603 | unknown | | |
| 371.358 | Chol | Cholesterol derivative[LM,d] | [M-H2O+H]+ | 645.561* | CE | CE(16:1)[a,b,c] | [M+Na]+ |
| 383.333 | oxChol | Cholesterol derivative[LM,d] | | 647.577* | CE | CE(16:0)[a,b,c] | [M+Na]+ |
| 385.348 | oxChol | Dehydrocholesterol[LM,d] | [M+H]+ | 652.610 | unknown | | |
| 401.343* | oxChol | 7-ketocholestol[d] | [M+H]+ | 654.628 | unknown | | |
| 428.370 | unknown | | | 656.630 | unknown | | |
| 429.375 | unknown | | | 666.495 | unknown | | |
| 430.379 | unknown | | | 668.525 | unknown | | |
| 431.385 | unknown | | | 668.622 | unknown | | |
| 496.346* | LPC | LPC(16:0)[d] | [M+H]+ | 669.563 | CE | CE(18:3)[a] | [M+Na]+ |
| 518.335* | LPC | LPC(18:3)[d] | [M+H]+ | 671.580* | CE | CE(18:2)[a,b,c] | [M+Na]+ |
| 520.351* | LPC | LPC(18:2)[d] | [M+H]+ | 673.593* | CE | CE(18:1)[a,b,c] | [M+Na]+ |
| 522.371* | LPC | LPC(18:1)[d] | [M+H]+ | 675.573 | SM | SM(d32:1)[a,b] | [M+H]+ |
| 524.374* | LPC | LPC(18:0)[d] | [M+H]+ | 676.560 | unknown | | |
| 542.322* | LPC | LPC(20:5)[d] | [M+H]+ | 678.623 | unknown | | |
| 544.340* | LPC | LPC(20:4)[d] | [M+H]+ | 680.636 | unknown | | |
| 546.351* | LPC | LPC(20:3)[d]<br>LPC(18:0)[b] | [M+H]+<br>[M+Na]+ | 681.643 | unknown | | |
| 549.488* | DAG | DAG(32:1)[a] | [M-H2O+H]+ | 682.653 | unknown | | |
| 551.509 | DAG | DAG(32:0)[a] | [M-H2O+H]+ | 683.658 | unknown | | |
| 575.510* | DAG | DAG(34:2)[a]<br>DAG(O-32:1)[b] | [M-H2O+H]+<br>[M+Na]+ | 685.567* | oxCE | Oxo-ODE-CE[d] | [M+Na]+ |
| 577.520* | DAG | DAG(34:1)[a]<br>DAG(O-32:0)[b] | [M-H2O+H]+<br>[M+Na]+ | 687.566* | oxCE*<br>CE | HODE-CE[b]<br>CE(18:2)[a] | [M+Na]+<br>[M+K]+ |
| 579.534 | DAG | DAG(34:0)[a] | [M-H2O+H]+ | 689.570 | CE<br>TAG | CE(18:1)[b]<br>TAG(38:1)[b] | [M+K]+<br>[M+Na]+ |
| 596.540 | unknown | | | 692.638 | unknown | | |
| 598.558 | unknown | | | 693.567* | CE | CE(20:5)[a,b] | [M+Na]+ |
| 599.497* | DAG | DAG(36:4)[a] | [M-H2O+H]+ | 695.581* | CE | CE(20:4)[a,b] | [M+Na]+ |
| 601.512* | DAG | DAG(36:3)[a]<br>DAG(O-34:2)[b] | [M-H2O+H]+<br>[M+Na]+ | 695.659 | unknown | | |
| 603.530* | DAG | DAG(36:2)[a]<br>DAG(O-34:1)[b] | [M-H2O+H]+<br>[M+Na]+ | 696.673 | unknown | | |
| 605.555 | DAG | DAG(36:1)[a] | [M-H2O+H]+ | 697.535* | SM | SM(32:1)[a,b] | [M+Na]+ |
| 610.572 | unknown | | | 697.591* | CE | CE(20:3)[a,b] | [M+Na]+ |
| 624.567 | unknown | | | 697.672 | unknown | | |
| 626.591 | unknown | | | 699.606 | CE | CE(20:2)[a,b] | [M+Na]+ |
| 627.544 | DAG | DAG(38:4)[a]<br>DAG(O-36:3)[b] | [M-H2O+H]+<br>[M+Na]+ | 701.571 | SM | SM(34:2)[b] | [M+H]+ |

*m/z = mass measured in MALDI-MSI experiment using Synapt G2Si TOF system. For exact mass measured with FTICR and ppm values, we refer to the online available METASPACE at https://metaspace2020.eu/ data named Human CEA Patient H - section 4 and Human CEA Patient I - section 4*

*lipid group = assigned lipid group based on database search, i.e. LipidMaps, HMDB.*

*ID = proven lipid identity, superscript denotes identification method: a. Lipidyzer MRM analysis, b. FTICR measurement combined with METASPACE database (FDR of maximum 10%), c. Identified in previous study [1], d. Identified from literature, LM: assigned from search in LipidMaps, HMDB: assigned from search in HMDB*

*adduct = positive ion adduct in identification experiment.*

** Asterisks denote the lipids that were included in the cross-correlation analysis*

[d] *Lipid identity not confirmed in identification experiments due to methodological limitations. However m/z values have been identified as 7-ketocholesterol [2,3], other cholesterol derivatives [3], LPCs [4] and oxo-ODE-CE [5] in literature and have been included as such in our cross-correlation analysis.*

Supplementary Table 1 – continued

| *m/z ± 0.02* | lipid class | ID | adduct | *m/z ± 0.02* | lipid class | ID | adduct |
|---|---|---|---|---|---|---|---|
| 703.583* | SM | SM(34:1)[a,b,c] | [M+H]+ | 792.600 | PC-O | PC(O-38:6) [LM,HMDB] | [M+H]+ |
| 705.589 | SM | SM(34:0)[b] | [M+H]+ | 794.614 | PC-O | PC(O-38:5) [LM,HMDB] | [M+H]+ |
| 707.542 | unknown | | | 796.541 | PC | PC(34:2)[a] | [M+K]+ |
| 711.550* | oxCE*<br>CE | CE(11:1D3)[HMDB]<br>CE(20:4)[a] | [M+Na]+<br>[M+K]+ | 796.615 | PC-O | PC(O-38:4) [LM,HMDB] | [M+H]+ |
| 717.578 | unknown | | | 798.548 | PC | PC(34:1)[a,b] | [M+K]+ |
| 719.571* | CE | CE(22:6)[a,b] | [M+Na]+ | 802.610 | unknown | | |
| 721.586* | CE | CE(22:5)[a,b,c] | [M+Na]+ | 803.617 | unknown | | |
| 723.544 | SM | SM(34:2)[b] | [M+Na]+ | 807.641* | SM | SM(40:2)[a,b] | [M+Na]+ |
| 725.565* | SM | SM(34:1)[a,b,c] | [M+Na]+ | 808.609 | PC | PC(36:2)[b] | [M+Na]+ |
| 727.566 | SM | SM(34:0)[b] | [M+Na]+ | 810.610* | PC | PC(38:4)[c] | [M+H]+ |
| 729.590* | SM | SM(36:2)[a,] | [M+H]+ | 811.676* | SM | SM(40:0)[b] | [M+Na]+ |
| 731.614* | SM | SM(36:1)[a,b,c] | [M+H]+ | 813.695* | SM | SM(42:2)[a,c] | [M+H]+ |
| 732.584 | unknown | | | 815.710 | SM | SM(42:1)[a,b] | [M+H]+ |
| 733.561 | unknown | | | 816.597 | PC-O | PC(O-38:5) [LM,HMDB] | [M+Na]+ |
| 734.580* | PC | PC(32:0)[a,b,c] | [M+H]+ | 820.539* | PC | PC(36:4)[a] | [M+K]+ |
| 735.572 | unknown | | | 822.555 | PC | PC(36:3)[a] | [M+K]+ |
| 736.577 | unknown | | | 823.592 | SM | SM(40:2)[a] | [M+K]+ |
| 739.550 | SM | SM(34:2)[LM,HMDB] | [M+K]+ | 824.584 | unknown | | |
| 741.545* | SM | SM(34:1)[a] | [M+K]+ | 825.610 | SM | SM(40:1)[a] | [M+K]+ |
| 743.555 | SM | SM(34:0)[LM,HMDB] | [M+K]+ | 827.622 | unknown | | |
| 746.606 | PC-O | PC(O-34:1) [LM,HMDB] | [M+H]+ | 827.709* | TAG | TAG(50:4)[a],<br>TAG(48:1)[a] | [M+H]+<br>[M+Na]+ |
| 750.565 | unknown | | | 828.620 | unknown | | |
| 751.574 | SM | SM(36:2)[a] | [M+Na]+ | 829.619 | unknown | | |
| 753.585 | SM | SM(36:1)[a,b] | [M+Na]+ | 829.717* | TAG | TAG(50:3)[a],<br>TAG(48:0)[a] | [M+H]+<br>[M+Na]+ |
| 755.552 | unknown | | | 830.574 | PC | PC(40:8)[b]<br>PC(38:5)[b] | [M+H]+<br>[M+Na]+ |
| 756.558* | PC | PC(32:0)[a,b] | [M+Na]+ | 832.600 | PC | PC(38:4)[b] | [M+Na]+ |
| 758.574* | PC | PC(34:2)[a,c] | [M+H]+ | 834.637 | unknown | | |
| 760.587* | PC | PC(34:1)[a,b,c] | [M+H]+ | 835.673* | SM | SM(44:5)[b]<br>SM(42:2) [a,b] | [M+H]+<br>[M+Na]+ |
| 762.618 | PC | PC(34:0)[HMDB] | [M+H]+ | 837.690 | SM | SM(42:1)[a] | [M+Na]+ |
| 764.652 | unknown | | | 839.590 | unknown | | |
| 766.578* | PC-O | PC(O-36:5) [LM,HMDB] | [M+H]+ | 846.661 | unknown | | |
| 766.754 | unknown | | | 847.593 | unknown | | |
| 768.581* | PC-O | PC(O-36:4) [LM,HMDB] | [M+H]+ | 848.588 | unknown | | |
| 772.542 | PC | PC(32:0)[a] | [M+K]+ | 848.676 | unknown | | |
| 774.580 | unknown | | | 849.611 | unknown | | |
| 776.584 | unknown | | | 850.605 | unknown | | |
| 780.564* | PC | PC(34:2)[a,b,c] | [M+Na]+ | 851.629 | SM | SM(42:2)[a] | [M+K]+ |
| 782.577* | PC | PC(36:4)[a,c]<br>PC(34:1)[a,b] | [M+H]+<br>[M+Na]+ | 853.721* | TAG | TAG(52:5)[a,b]<br>TAG(50:2)[a,] | [M+H]+<br>[M+Na]+ |
| 784.591* | PC | PC(36:3)[a,b,c] | [M+H]+ | 855.592 | unknown | | |
| 786.610* | PC | PC(36:2)[a,b,c] | [M+H]+ | 855.737 | TAG | TAG(52:4),<br>TAG(50:1) [LM,HMDB] | [M+H]+<br>[M+Na]+ |
| 788.620* | PC | PC(36:1)[c] | [M+H]+ | 856.596 | PC | PC(42:9)<br>PC(40:6)[HMDB] | [M+H]+<br>[M+Na]+ |
| 790.585 | unknown | | | 857.596 | unknown | | |

Supplementary Table 1 – continued

| *m/z ± 0.02* | lipid group | ID | adduct |
|---|---|---|---|
| 857.750* | TAG | TAG(52:3)[a], TAG(50:0)[a] | [M+H]+ [M+Na]+ |
| 860.774 | unknown | | |
| 862.790 | unknown | | |
| 870.684 | PC | PC(42:2) [HMDB] | [M+H]+ |
| 872.693 | PC | PC(42:1) [HMDB] | [M+H]+ |
| 877.738* | TAG | TAG(54:7)[a], TAG(52:4)[a] | [M+H]+ [M+Na]+ |
| 879.738* | TAG | TAG(54:6) [a], TAG(52:3)[a] | [M+H]+ [M+Na]+ |
| 881.760* | TAG | TAG(54:5)[a], TAG(52:2)[a] | [M+H]+ [M+Na]+ |
| 883.768* | TAG | TAG(54:4), TAG(52:1)[a] | [M+H]+ [M+Na]+ |
| 885.781* | TAG | TAG(54:3), TAG(52:0)[a] | [M+H]+ [M+Na]+ |
| 888.804 | unknown | | |
| 889.726 | unknown | | |
| 895.725 | unknown | | |
| 897.736 | PE-Cer | PE-Cer(t38:0)[b] | [M+Na]+ |
| 901.569 | unknown | | |
| 901.732* | TAG | TAG(56:9), TAG(54:6)[a] | [M+H]+ [M+Na]+ |
| 903.751* | TAG | TAG(56:8), TAG(54:5)[a] | [M+H]+ [M+Na]+ |
| 905.751* | TAG | TAG(56:7), TAG(54:4)[a] | [M+H]+ [M+Na]+ |
| 907.773* | TAG | TAG(56:6), TAG(54:3)[a] | [M+H]+ [M+Na]+ |
| 909.794* | TAG | TAG(56:5), TAG(54:2)[a] | [M+H]+ [M+Na]+ |
| 923.750 | unknown | | |
| 927.744* | TAG | TAG(58:10), TAG(56:7)[a] | [M+H]+ [M+Na]+ |
| 929.768* | TAG | TAG(58:9), TAG(56:6)[a] | [M+H]+ [M+Na]+ |
| 931.778* | TAG | TAG(58:8), TAG(56:5)[a] | [M+H]+ [M+Na]+ |
| 932.570 | unknown | | |
| 933.790* | TAG | TAG(58:7), TAG(56:4)[a] | [M+H]+ [M+Na]+ |
| 940.770 | unknown | | |
| 955.774* | TAG | TAG(60:10), TAG(58:7)[a] | [M+H]+ [M+Na]+ |
| 956.570 | unknown | | |
| 958.583 | unknown | | |
| 980.570 | unknown | | |
| 984.602 | unknown | | |
| 986.769 | unknown | | |
| 1,010.774 | unknown | | |
| 1,012.783 | unknown | | |
| 1,014.798 | unknown | | |
| 1,034.772 | unknown | | |
| 1,036.783 | unknown | | |
| 1,038.805 | unknown | | |
| 1,040.817 | unknown | | |
| 1,042.824 | unknown | | |
| 1,077.583 | unknown | | |

Supplementary Figure 1: Histological tissue composition of 12 carotid plaques

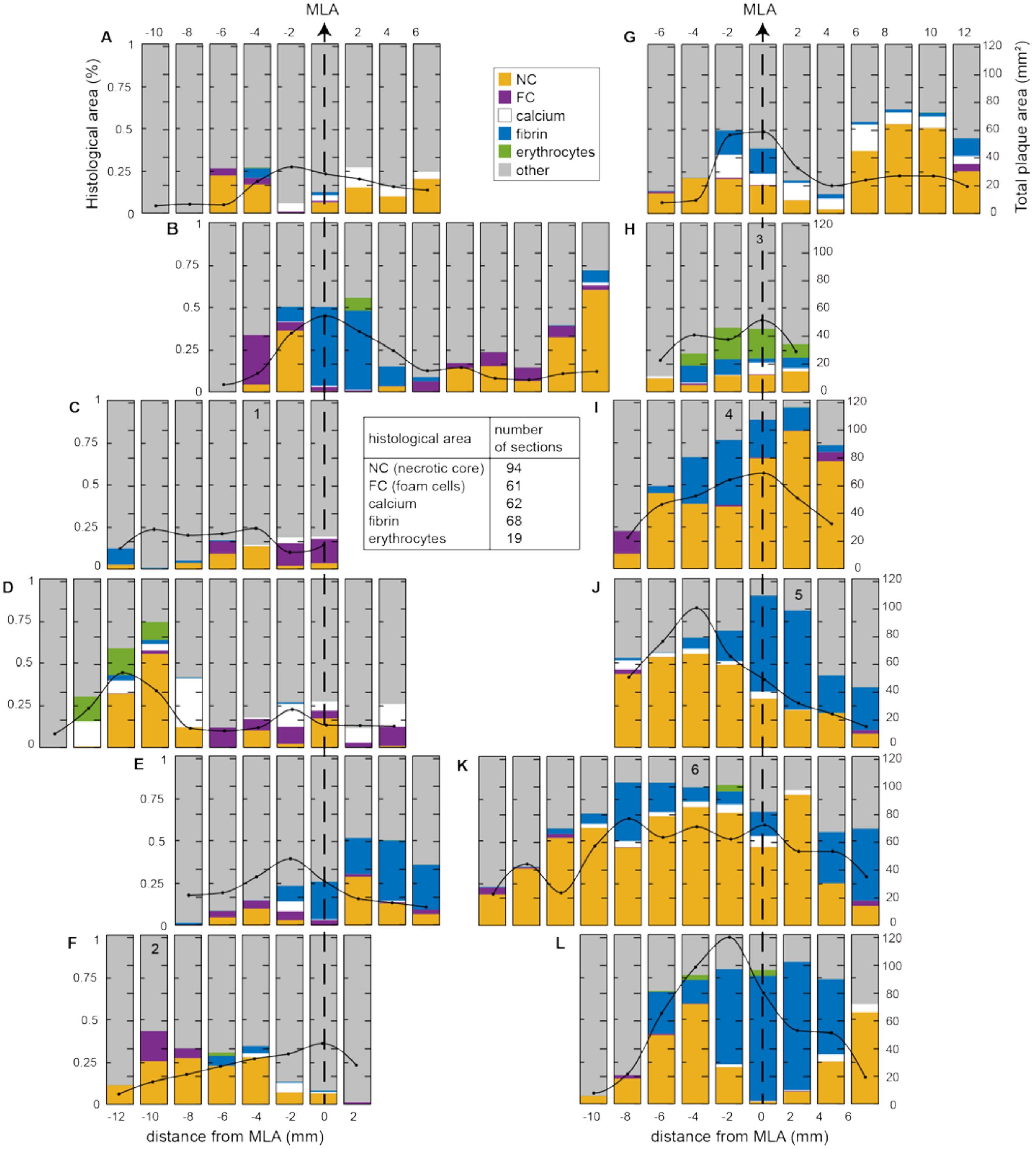


| histological area | number of sections |
|---|---|
| NC (necrotic core) | 94 |
| FC (foam cells) | 61 |
| calcium | 62 |
| fibrin | 68 |
| erythrocytes | 19 |

*Supplementary Figure 1: Overview of plaque components present in each carotid endarterectomy (CEA) sample. A-L: Bar graphs show the relative proportions (%) of necrotic core (NC), foam cells (FC), calcifications, fibrin and erythrocytes compared to the total intima area, see left axes.Tissue labeled as other was not segmented, this part of the tissue was histologically heterogeneous, but did not fit into any accepted classifications of atherosclerotic tissues. Total intima area represents the area of the intima layer of the vessel. The black line superimposed on the bargraph shows the total intima area in mm$^2$, see right axis. The numbers in the graphs correspond to the exemplatory sections shown in Figure 1, 2 and 4. The table in the middle shows the frequency of occurance of each histological component. MLA = minimal lumen area. Per CEA sample, the tissue section containing the minimal lumen area is denoted.*

Supplementary Table 2: OPLS-DA model parameters

| **Histological area** | **$R^2$** | **$Q^2$** | **CV-ANOVA** | **Patients included in model** *(number of patients in model / total number of patients)* ***** |
|---|---|---|---|---|
| NC | 0.57 | 0.51 | 5.34E-19 | All except D, H, L *(9/12)* |
| Fibrin | 0.68 | 0.52 | 2.53E-07 | B, C, F, G, J, K *(6/12)* |
| Foam cells | 0.63 | 0.55 | 7.33E-15 | All except D *(11/12)* |
| Erythrocytes | 0.67 | 0.61 | 5.57E-05 | B, F, H, I, K, L *(6/12)* |

*This table summarizes the parameters for OPLS-DA models comparing the mean spectrum of a histological area (NC, fibrin, FCs or erythrocytes) to the mean spectrum of tissue outside this histological area. The patients included in the model are reported in the last column of this table, all tissue sections of included patients were added to the model. $R^2$ and $Q^2$ values represent the quality of fit and predictability of the model respectively. Significance of OPLS-DA models was checked by sevenfold cross-validation analysis of variance (CV-ANOVA).*

** The number of patients for which $R^2$ and $Q^2$ values were highest. Minimal 6 out of 12 patients were included in the model.*

Supplementary Table 3: Lists of *m/z* values with VIP>1.0 resulting from significant OPLS-DA models

**Necrotic core**

*M/z* values with VIP>1.0, resulting from the OPLS-DA model comparing NC with not-NC areas

| ***m/z* value** | **VIP** |
|---|---|
| 811.676 | 2.00 |
| 701.571 | 1.98 |
| 729.590 | 1.95 |
| 685.567 | 1.94 |
| 839.590 | 1.93 |
| 675.573 | 1.92 |
| 689.570 | 1.91 |
| 813.694 | 1.86 |
| 705.589 | 1.85 |
| 703.583 | 1.84 |
| 717.578 | 1.80 |
| 857.596 | 1.79 |
| 687.565 | 1.78 |
| 676.560 | 1.77 |
| 855.592 | 1.77 |
| 815.710 | 1.76 |
| 721.585 | 1.73 |
| 711.550 | 1.72 |
| 834.637 | 1.72 |
| 719.571 | 1.71 |
| 522.371 | 1.69 |
| 723.544 | 1.68 |
| 727.566 | 1.63 |
| 697.535 | 1.62 |
| 856.596 | 1.61 |
| 835.673 | 1.59 |
| 671.580 | 1.58 |
| 673.593 | 1.58 |
| 544.340 | 1.54 |
| 731.614 | 1.53 |
| 746.606 | 1.50 |
| 796.615 | 1.50 |
| 645.561 | 1.50 |
| 520.351 | 1.47 |
| 794.614 | 1.47 |

| ***m/z* value** | **VIP** |
|---|---|
| 743.555 | 1.45 |
| 849.610 | 1.44 |
| 647.577 | 1.42 |
| 901.568 | 1.41 |
| 823.592 | 1.41 |
| 739.549 | 1.41 |
| 751.574 | 1.41 |
| 837.690 | 1.39 |
| 766.578 | 1.39 |
| 524.374 | 1.37 |
| 725.565 | 1.36 |
| 669.563 | 1.33 |
| 825.610 | 1.32 |
| 496.346 | 1.31 |
| 807.641 | 1.31 |
| 768.581 | 1.30 |
| 542.322 | 1.29 |
| 792.600 | 1.28 |
| 851.629 | 1.26 |
| 850.605 | 1.23 |
| 695.581 | 1.21 |
| 816.597 | 1.20 |
| 695.659 | 1.18 |
| 753.585 | 1.15 |
| 546.351 | 1.14 |
| 847.593 | 1.14 |
| 696.673 | 1.12 |
| 790.585 | 1.12 |
| 518.335 | 1.11 |
| 697.672 | 1.07 |
| 1,077.583 | 1.06 |
| 692.638 | 1.05 |
| 827.622 | 1.05 |
| 385.347 | 1.01 |

**Fibrin**

*M/z* values with VIP>1.0, resulting from the OPLS-DA model comparing fibrin with not-fibrin areas

| ***m/z* value** | **VIP** |
|---|---|
| 931.778 | 2.12 |
| 929.768 | 2.06 |
| 955.774 | 2.05 |
| 383.333 | 2.01 |
| 927.744 | 1.86 |
| 599.497 | 1.84 |
| 401.343 | 1.84 |
| 905.751 | 1.81 |
| 901.732 | 1.78 |
| 603.529 | 1.78 |
| 933.790 | 1.78 |
| 601.512 | 1.74 |
| 721.585 | 1.71 |
| 923.750 | 1.68 |
| 909.794 | 1.66 |
| 907.773 | 1.63 |
| 903.751 | 1.62 |
| 605.555 | 1.62 |
| 829.717 | 1.62 |
| 579.534 | 1.61 |
| 575.510 | 1.61 |
| 857.596 | 1.59 |
| 551.509 | 1.55 |
| 856.596 | 1.52 |
| 792.600 | 1.51 |
| 857.750 | 1.51 |
| 549.488 | 1.46 |
| 577.520 | 1.45 |
| 666.495 | 1.44 |
| 853.721 | 1.43 |
| 855.737 | 1.42 |

| ***m/z* value** | **VIP** |
|---|---|
| 746.606 | 1.42 |
| 889.726 | 1.39 |
| 796.615 | 1.37 |
| 895.725 | 1.36 |
| 877.738 | 1.34 |
| 794.614 | 1.32 |
| 719.571 | 1.29 |
| 676.560 | 1.26 |
| 827.709 | 1.25 |
| 627.544 | 1.23 |
| 766.578 | 1.22 |
| 1,040.817 | 1.21 |
| 883.768 | 1.20 |
| 733.561 | 1.19 |
| 879.738 | 1.17 |
| 685.567 | 1.15 |
| 855.592 | 1.13 |
| 701.571 | 1.13 |
| 816.597 | 1.12 |
| 1,038.805 | 1.12 |
| 897.736 | 1.11 |
| 762.618 | 1.11 |
| 1,036.783 | 1.11 |
| 729.590 | 1.08 |
| 839.590 | 1.07 |
| 1,042.824 | 1.06 |
| 385.347 | 1.04 |
| 705.589 | 1.03 |
| 881.760 | 1.02 |
| 885.781 | 1.02 |
| 692.638 | 1.00 |

**Foam cells**

*M/z* values with VIP>1.0, resulting from the OPLS-DA model comparing foam cell with not-foam cell areas

| ***m/z* value** | **VIP** |
|---|---|
| 697.591 | 1.52 |
| 888.804 | 1.47 |
| 729.590 | 1.42 |
| 862.790 | 1.41 |
| 1,010.774 | 1.41 |
| 685.567 | 1.40 |
| 986.769 | 1.38 |
| 756.558 | 1.35 |
| 780.564 | 1.33 |
| 848.588 | 1.32 |
| 885.781 | 1.30 |
| 782.577 | 1.28 |
| 827.709 | 1.28 |
| 638.472 | 1.26 |
| 1012.78 | 1.26 |
| 784.591 | 1.26 |
| 732.584 | 1.24 |
| 850.605 | 1.24 |
| 829.717 | 1.24 |
| 755.552 | 1.22 |
| 808.609 | 1.22 |
| 860.774 | 1.22 |
| 758.574 | 1.22 |
| 734.580 | 1.21 |
| 772.542 | 1.18 |
| 627.544 | 1.17 |
| 824.584 | 1.17 |
| 848.676 | 1.16 |
| 786.610 | 1.16 |
| 760.587 | 1.15 |

| ***m/z* value** | **VIP** |
|---|---|
| 1,042.824 | 1.15 |
| 1,038.805 | 1.15 |
| 697.672 | 1.14 |
| 1,036.783 | 1.14 |
| 849.610 | 1.13 |
| 830.574 | 1.13 |
| 822.555 | 1.13 |
| 764.652 | 1.12 |
| 731.614 | 1.12 |
| 870.684 | 1.12 |
| 851.629 | 1.12 |
| 798.548 | 1.12 |
| 825.610 | 1.11 |
| 832.600 | 1.11 |
| 673.593 | 1.11 |
| 1,034.772 | 1.10 |
| 735.572 | 1.10 |
| 796.541 | 1.07 |
| 827.622 | 1.07 |
| 810.610 | 1.05 |
| 733.561 | 1.05 |
| 701.571 | 1.05 |
| 1,014.798 | 1.04 |
| 855.592 | 1.04 |
| 839.590 | 1.04 |
| 932.570 | 1.04 |
| 676.560 | 1.02 |
| 846.661 | 1.01 |
| 687.565 | 1.00 |

**Erythrocytes**

*M/z* values with VIP>1.0, resulting from the OPLS-DA model comparing erythrocyte with not-erythrocyte areas

| ***m/z* value** | **VIP** |
|---|---|
| 932.570 | 1.35 |
| 780.564 | 1.35 |
| 782.577 | 1.34 |
| 958.583 | 1.33 |
| 756.558 | 1.32 |
| 758.574 | 1.32 |
| 786.610 | 1.31 |
| 784.591 | 1.31 |
| 768.581 | 1.31 |
| 760.587 | 1.30 |
| 832.600 | 1.29 |
| 956.569 | 1.29 |
| 810.610 | 1.29 |
| 790.585 | 1.28 |
| 734.580 | 1.28 |
| 796.541 | 1.28 |
| 830.574 | 1.26 |
| 808.609 | 1.26 |
| 772.542 | 1.25 |
| 735.572 | 1.24 |
| 798.548 | 1.24 |
| 820.539 | 1.22 |
| 984.602 | 1.21 |
| 822.555 | 1.21 |
| 788.620 | 1.20 |
| 905.751 | 1.19 |
| 980.570 | 1.17 |
| 766.578 | 1.15 |
| 736.577 | 1.14 |
| 903.751 | 1.11 |
| 923.750 | 1.11 |
| 762.618 | 1.11 |
| 707.542 | 1.10 |
| 796.615 | 1.08 |
| 897.736 | 1.07 |
| 816.597 | 1.07 |
| 929.768 | 1.07 |
| 755.552 | 1.05 |
| 877.738 | 1.03 |
| 931.778 | 1.03 |
| 955.774 | 1.02 |
| 652.609 | 1.01 |

Supplementary Table 4: Number of *m/z* values with VIP > 1.0 for the different multivariate models and the NMF component in which these *m/z* values are most abundant

| **NMF component** | **NC** | **Fibrin** | **FC** | **Erythrocytes** |
|---|---|---|---|---|
| 1 | 18 | 4 | 9 | 0 |
| 2 | 2 | 4 | 0 | 0 |
| 3 | 0 | 21 | 2 | 10 |
| 4 | 2 | 0 | 0 | 0 |
| 5 | 39 | 13 | 1 | 0 |
| 6 | 22 | 9 | 21 | 29 |
